# CytoSpatio: Learning cell type spatial relationships using multirange, multitype point process models

**DOI:** 10.1101/2024.10.31.621408

**Authors:** Haoran Chen, Robert F. Murphy

**Affiliations:** Computational Biology Department School of Computer Science, Carnegie Mellon University

**Keywords:** point process models, generative models, cell types, tissue images, spatial relationships, multiplexed fluorescence imaging, synthetic tissue simulation, spatial proteomics

## Abstract

Recent advances in multiplexed fluorescence imaging have provided new opportunities for deciphering the complex spatial relationships among various cell types across diverse tissues. We introduce CytoSpatio, open-source software that constructs generative, multirange, and multitype point process models that capture interactions among multiple cell types at various distances simultaneously. On analyzing five cell types across five tissues, our software showed consistent spatial relationships within the same tissue type, with certain cell types like proliferating T cells consistently clustering across tissue types. It also revealed that the attraction-repulsion relationships between cell types like B cells and CD4-positive T cells vary with tissue type. CytoSpatio can also generate synthetic tissue structures that preserve the spatial relationships seen in training images, a capability not provided by previous descriptive, motif-based approaches. This potentially allows spatially realistic simulations of how cell relationships affect tissue biochemistry.

## Introduction

The functions of a tissue are often determined by the type and arrangement of its constituent cells. Distinct shapes, sizes, and molecular properties of cell types lead to specialized functions within a tissue [1-5]. However, spatial relationships among various cell types within diverse tissues are often more complex, and their impact on tissue functions is not fully understood.

Traditional imaging techniques, such as confocal microscopy, electron microscopy, and computed tomography (CT), have allowed scientists to investigate the spatial relationships between specific cell types within particular tissues [6-9]. However, these approaches typically required manual annotations of cell types. Therefore, they faced challenges of subjectivity in cell type annotations, limited scalability of conclusions across tissues, and most notably, the inability to capture complex spatial relationships due to the restriction on the number of identifiable cell types.

Recent advances in multiplexed imaging approaches for spatial transcriptomics and proteomics offer an unprecedented opportunity for researchers to explore the spatial relationships between a diverse range of cell types simultaneously [10-14]. By employing biomarkers targeting distinct RNA transcripts or proteins within cells in a multiplexed manner, various cell types can be concurrently visualized in tissue samples [15-17].

This advancement has motivated researchers to investigate spatial relationships among cell types with a variety of methods, mainly involving quantification and summarization of colocalization and correlation between cell types using analytic and statistical methods.

Behanova et al. [18] summarized and reviewed a variety of spatial statistics methods, tools, and software. The primary focus was on testing various hypotheses regarding whether cell types are randomly distributed, rather than attempting to construct models to capture complex spatial relationships.

Filipek et al. [19] presented CytoMAP, a spatial analysis platform that quantified local cell composition and global tissue structure. This platform defines cell-centered local neighborhoods across the tissue, and groups similar neighborhoods together through clustering methods. It provides overall correlation and neighborhood composition between cell types for colorectal tumor and lymphoid tissues. While CytoMAP is a powerful tool for the spatial analysis of cell type relationships in tissue images, it has certain limitations. First, choice of the range for cell-centered local neighborhoods would be expected to significantly affect results. This could limit the reproducibility of the analyses and the comparability of spatial relationships between cell types in different tissues. Second, while CytoMAP does offer correlations and neighborhood compositions between cell types, it may oversimplify the complexity of spatial relationships among various cell types, a common concern shared with spatial statistics.

Barlow et al. [20] hypothesized that tissues are composed hierarchically from smaller to larger components following certain assembly rules. To test this hypothesis, a hierarchical computational framework was devised to systematically identify the characteristic local compositions of cell types, known as cellular neighborhoods, map the local interactions and co-localization of these neighborhoods into distinct microenvironments, and delineate assembly rules that govern the formation of these microenvironments into tissue motifs. This hierarchical analysis produced proposed assembly rules for normal lymph node, spleen, and tonsil tissue, as well as colorectal cancer tissue. However, like CytoMAP, both the specific choices of the hierarchical design and the fixed parameters used to define the ranges of neighborhoods and microenvironments were not well justified or explored. The approach was also not incorporated into a probabilistic, generative framework to allow estimation of the likelihood of a tissue image being produced by a given model and/or the quantitative similarity between different tissues, and to allow generation of synthetic tissue images.

To address the limitations of these existing methods, we sought to employ generative statistical models to learn and represent the complex spatial relationships between different cell types in different tissues beyond pairwise analyses of colocalization and correlation.

Spatial point process models [21] are generative statistical models designed to learn the probability of individual objects (points) occurring at specific locations in space, including dependence of that probability on locations of other objects. The collection of points (including their locations within a defined region) are referred to as a “point pattern”, and models capturing how such point patterns are generated are referred to as “point process models.” These models have found widespread application in the analysis of spatial relationships across various domains, such as meteorology [22], ecology [23], criminology [24], and social sciences [25]. In cell biology, spatial point process models have been employed to elucidate the spatial relationships between punctate organelles and various cellular components, such as the nuclear membrane and microtubules [26, 27]. They have also been used to investigate the assembly of viral ribonucleoprotein complexes [28] and to identify prognostic structural features in colon cancer tissues [29]. Although these point process models have been successful in revealing spatial dependencies and interaction patterns between objects in different contexts, they typically focus on one type of object at a time. In these models, the locations of other point types, if they exist, are treated as influential “factors” that may affect the spatial distribution of the target point type. Consequently, separate models must be trained for each object type. To address this limitation, we chose to implement a point process model capable of simultaneously learning the spatial relationships between many types of objects, namely multitype point process model (or marked point process model) [30-34].

In a multitype point process model, when assuming there are interactions between different types of points, a common challenge is to determine the maximum interaction distance over which two types of points can influence each other. Conventionally, a range parameter was determined either by the distance from the nearest neighbor up to the distance of commonly observed interactions between two types of points [35] or by a distance distribution of nearest-neighbor between two types of points [36]. While these approaches offer a useful approximation, they are highly dependent on empirical observations from the data set. If the data set is limited or biased, they might lead to an inaccurate estimation of the maximum interaction distance. They might also constrain the extent to which models trained on different tissues may be compared. To overcome this challenge, we designed multirange models wherein different types of points can influence each other differently based on a specific range. This allows greater sensitivity in distinguishing different types of interactions.

In this study, we introduce CytoSpatio, open-source software that constructs generative, multirange, multitype point process models. We demonstrate its superior performance over single-range models using images from five different tissues containing five distinct cell types. We show how the models can be used to compare cell type spatial relationships between images from the same tissue or between images of different tissues. Additionally, we can use our approach to evaluate heterogeneity in different tissue subregions. Perhaps most usefully, we construct interaction network graphs that directly exhibit and compare the spatial relationships among cell types. Lastly, we demonstrate that our models can generate synthetic tissue images that preserve the spatial relationships observed in real tissue images, enabling rigorous validation of the models. Figure 1 illustrates the processes involved in constructing models using our approach.

**Figure 1.**
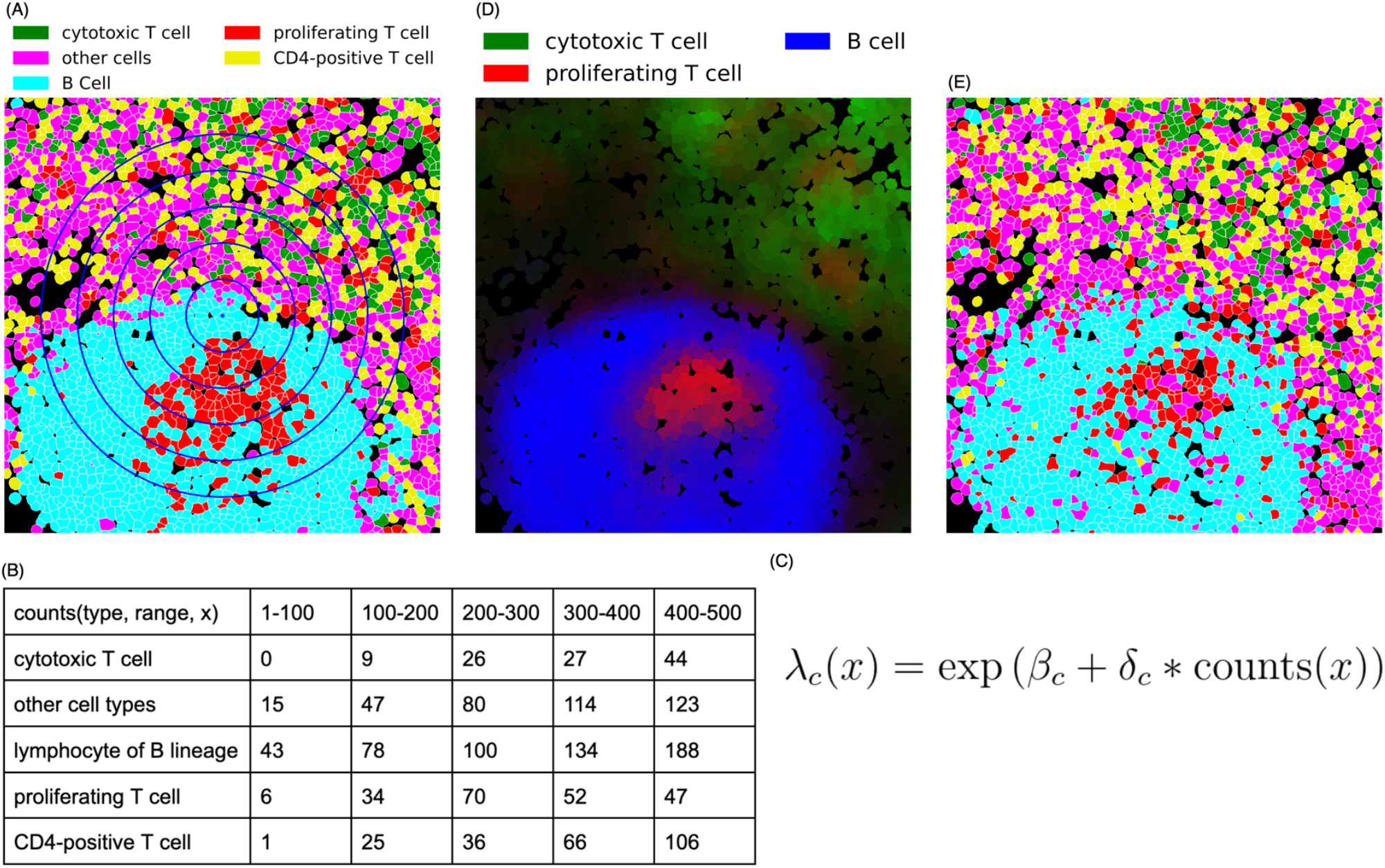
CytoSpatio process for learning spatial relationships between different cell types. (A) A region from a larger lymph node image is shown, with cell types shown in different colors and cell boundaries shown in white. The blue concentric circles denote five distance ranges of 100-500 pixels at 100-pixel intervals. (B) The training process involves counting the number of other cells of each type within varying distance ranges for each cell, as illustrated for the central cell (small blue diamond) in panel A, a B cell. (C) A simplified version of the equation used for the fitting process in a point process model to learn the spatial relationships among cell types is shown. The probability λ of a particular cell type *c* at a given location, x, is given by a (global) base intensity (β) adjusted for the influence of (multiplied by) the local frequencies of all cell types. This adjustment is given by the dot product of a vector of interaction coefficients (*δ*) for this cell type with all cell types (including its own) and a vector (Counts(x)) reflecting the counts of each cell type. The interaction coefficient and counts can be for a single range (i.e., one of the columns in panel B) or can be concatenated across multiple ranges (i.e., linearizing the counts in panel B). (D) Predicted intensities (proportional to the probabilities of occurrence) are shown for three cell types for each cell in this region (derived from a model trained with the entire image). Brighter colors indicate a higher predicted intensity, with each color corresponding to a distinct cell type. (E) A synthetic image depicting predicted cell types generated for this region from the model is shown. The image was generated from the model using the positions of each cell in panel A but assigning each cell’s type based on the predicted probabilities across the cell types for that location (cell type colors are the same as in panel (A)).

## Results

For this study, we used multiplexed tissue images from the Human BioMolecular Atlas Program (HuBMAP) [37]. Images for five tissues were segmented into single cells and the cell type of each cell was assigned as described in the Methods.

### Assessing non-randomness of cell type distributions in different tissues

We began our analysis by exploring whether the cell type distribution in each tissue is random, which would imply a lack of meaningful spatial relationships among cell types. We posed a null hypothesis that the cell type distribution in a tissue image would be equivalent to a distribution with the same cell locations but randomized cell types. For each tissue, we randomized the cell types within all images 100 times, generating 100 sets of point patterns with shuffled cell types. These patterns served as a background distribution for our hypothesis testing. For each set, we trained a multitype Strauss Hardcore model (see **Methods**) with the range that two cell types can affect each other (referred to as a Straus radius) set to 100 pixels and the range within which two cells cannot come closer to each other (referred to as a Hardcore radius) set to 1 pixel (1 pixel equals 0.377 microns). To measure agreement between a model and a set of point patterns, we used a metric that quantified the average disparity between each point pattern and the predicted intensity from the model (average deviance per cell, see **Methods**). For each shuffled model, we measured average deviance per cell against a randomly selected shuffled point pattern set from the same tissue, and also against the unshuffled point pattern from the original image.

As shown in Figure 2, we consistently observed that the average deviance per cell was lower when the models trained on a shuffled pattern set were tested against another shuffled point pattern set (red boxplots), as compared to when tested on the original point pattern set (green boxplots). As expected, this indicates for each tissue that the shuffled pattern sets were more similar to each other than they were to the original (unshuffled) pattern set. This strong deviation of the original patterns from randomness was statistically significant (p<0.01) for all five tissue types investigated. Interestingly, we found that the cell type distributions in thymus, small intestine (SI), and large intestine (LI) were particularly non-random, resulting in significantly higher deviance when their randomized models were tested against the original patterns.

**Figure 2.**
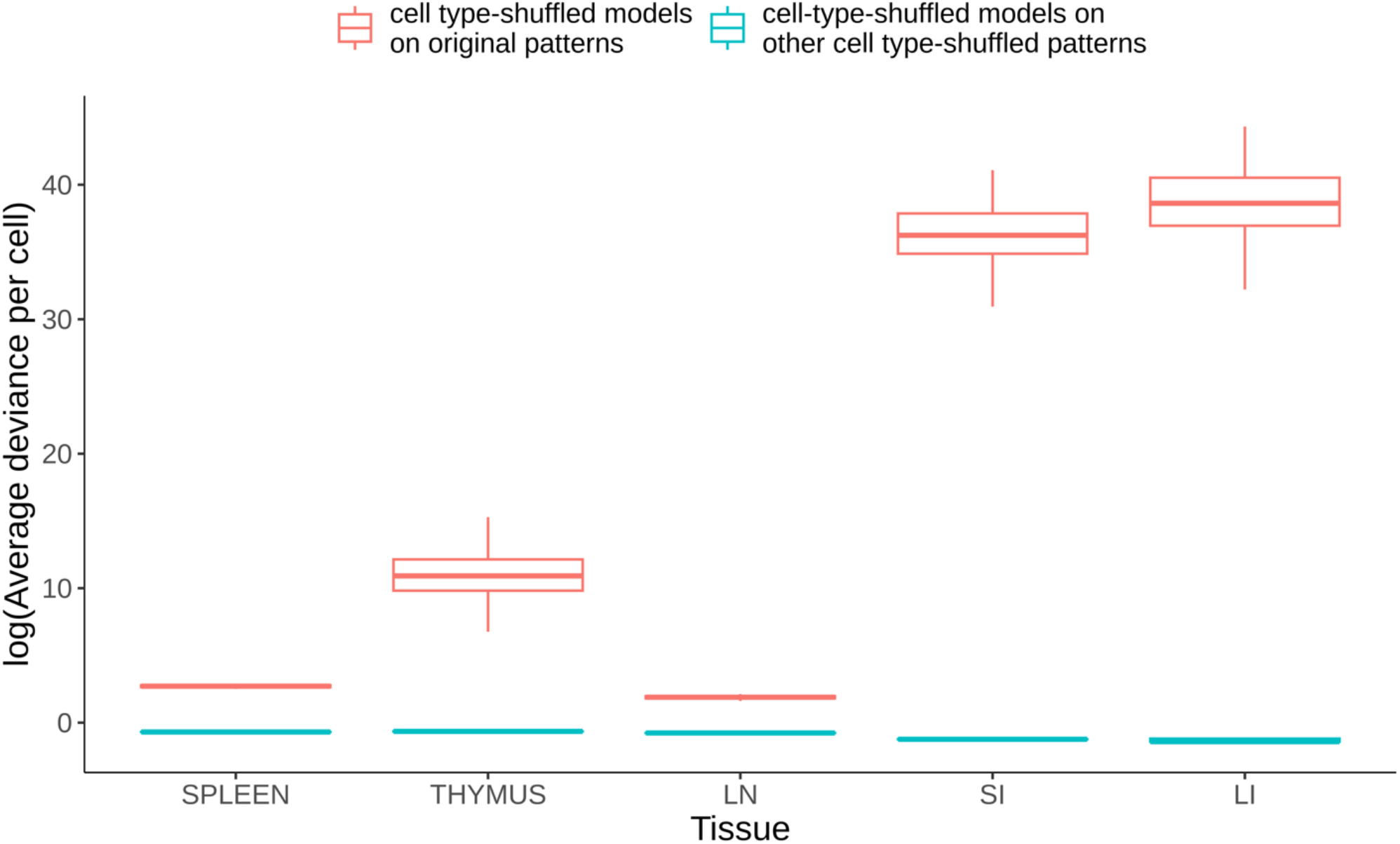
Comparison of average deviance per cell between shuffled point pattern sets and original point pattern sets. Lower average deviance per cell indicates a higher likelihood that a particular image could have been produced by a given model. The average deviance per cell is depicted in the boxplots, with the red boxplots representing the deviances when models trained on a shuffled point pattern set were compared to another shuffled point pattern set. The green boxplots represent the deviances when the same models were compared to the original point pattern set. Whiskers are drawn at 1.5 times the difference between the first and third quartiles. The significantly higher deviances for the original patterns compared to those for the shuffled patterns demonstrate the non-random distribution of cell types within the tissues studied. How the extremely high deviances seen in some cases can be obtained is discussed in the **Methods**.

### Comparing multirange to single range of Strauss Hardcore

We next evaluated whether our multirange, multitype Strauss-Hardcore model (see **Methods**) provides a more accurate fit for learning spatial relationships among cell types in our tissue images, compared to conventional Strauss Hardcore models with a single Strauss radius. For each tissue, we trained Strauss Hardcore models using various single radii (in pixels), as well as our multirange model that incorporates five distinct Strauss radii ranging from 100 to 500 at 100-pixel intervals.

An important component of constructing point process models is the creation of “dummy” points that have different types than the observed points so that the model can learn not only that observed points should have high probability, but that non-observed point should in general have low probability (see **Methods**). In order to compare models for different radii, we evaluated each model’s goodness-of-fit using the average deviance per real cell, per dummy cell, and per both real and dummy cells.

Figure 3 shows that, compared to the conventional Strauss Hardcore models with five single ranges, our multirange model consistently yielded the lowest average deviances for all five tissue types. Interestingly, we observed a gradual decline in the performance of the single radius model as the Strauss radius expanded from 100 to 500 pixels. This implies that the positioning of specific cell types is primarily influenced by their proximate neighboring cells, while cells at greater distances may introduce mixed spatial relationships that lower the prediction accuracy. Despite this, the spatial information derived from cells at larger distances remains beneficial for predicting cell types, contributing to the superior accuracy of the multi-range model across the five tissue types.

**Figure 3.**
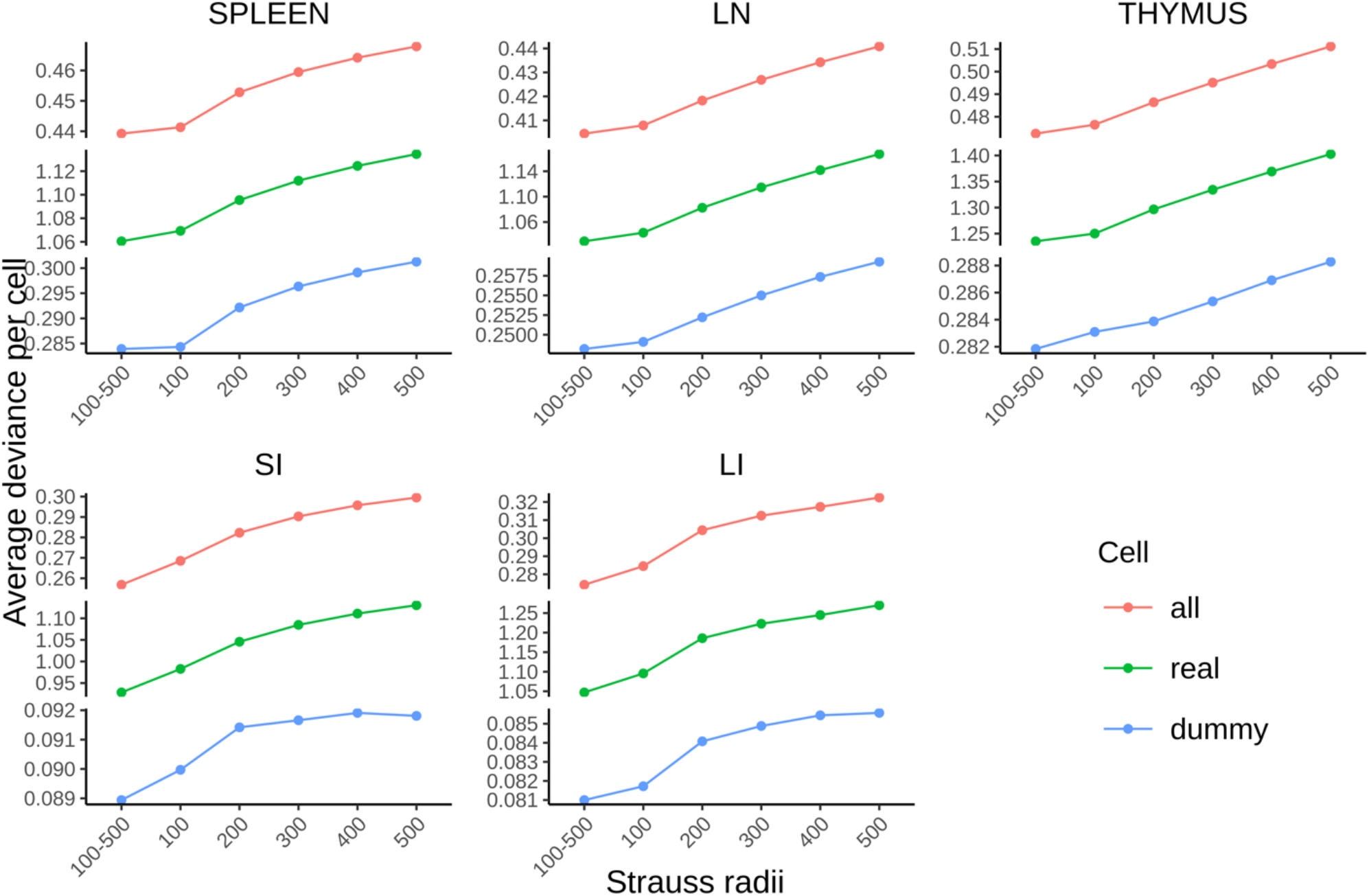
Performance comparison between multirange and single range multitype Strauss Hardcore models. The average deviance per cell for all cells, real cells, and dummy cells respectively measure the overall goodness-of-fit of the model, the prediction accuracy of cell types at their locations, and the accuracy of predicting locations devoid of real cells.

It is important to consider the relationships between the radius ranges used in constructing the models, the radii of the cell types being considered, and the size of the image pixels. For images with the same pixel size and similar cell radii, models can be directly compared (as we have here). As long as pixel size of the image (the width and height of each pixel in the sample plane; 0.377 microns for the images analyzed here) is reasonably smaller than the typical radii of the cell types, it does not significantly affect the estimation of cell-cell distances (when expressed in microns). Models for images of different pixel sizes can also be compared as long as the radius ranges (in pixels) are adjusted for each image so that they represent the same length (in microns).

### Evaluating differences in cell type spatial relationships within and across tissues

We next asked, using two distinct approaches, whether spatial relationships among cell types were more similar within the same tissue than they were between different tissues. Both approaches used sets of models for each tissue that were derived from a leave-one-out cross-validation process (see **Methods**).

The first approach involved calculating the Gaussian kernel similarity between the concatenated vectors of interaction coefficients for all radii (which encode the attraction or repulsion among cell types) of a pair of models. To provide an overall measure of similarity within or between tissues, we averaged similarity values between all pairs of models for a given tissue, and between all pairs of models from two tissues (Figure 4A). We found that models trained on images from the same tissue yielded similarities very close to 1. The value for lymph node were slightly lower, indicating slightly more heterogeneity among the images of that tissue. All of the same tissue values were consistently higher than those comparing models from different tissues. However, spleen, lymph node, and thymus tissues were more similar to each other than any of them were to either large or small intestine (which we quite similar to each other).

**Figure 4.**
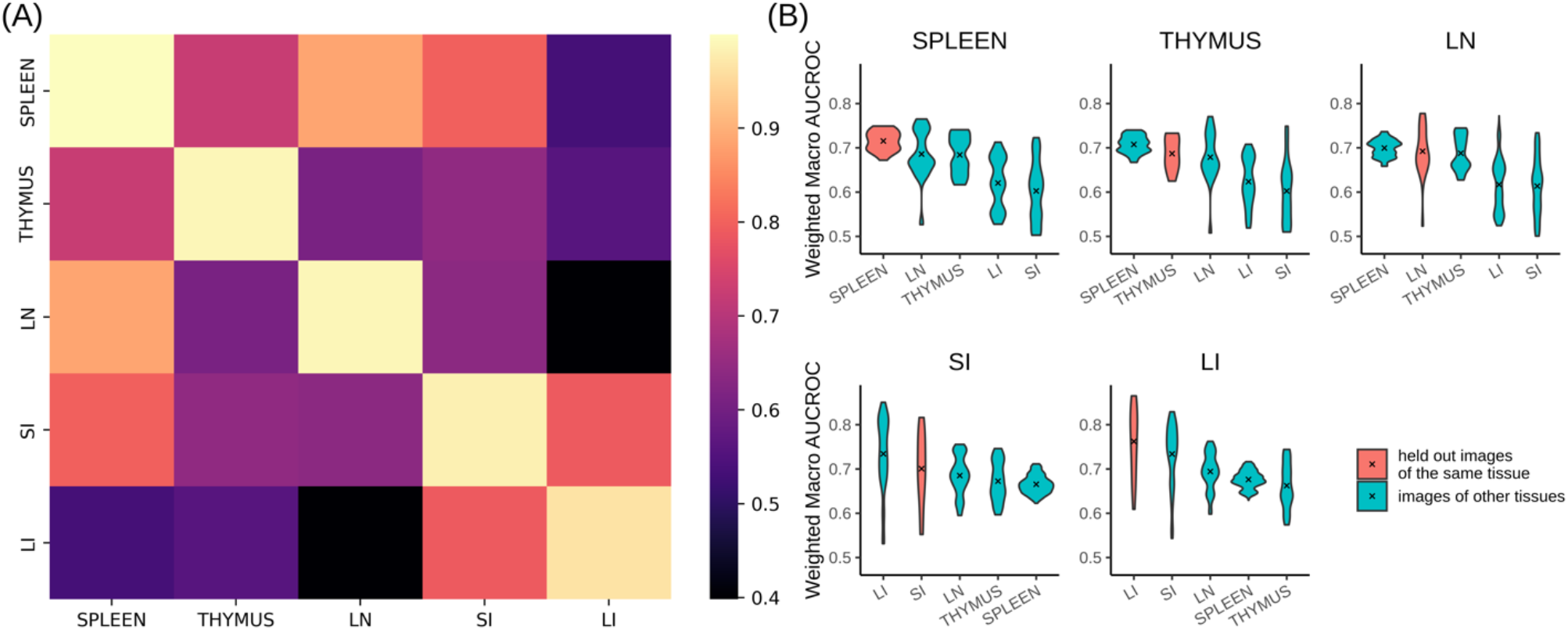
Comparison of cell type spatial relationships within and across different tissues. (A) The interaction coefficients between models are directly compared using Gaussian kernel similarity. Lighter color indicates greater similarity. (B) The predictive accuracy on held-out images of a given tissue as well as images from other tissues was measured using wmAUC. In each tissue panel, the violin plots are arranged in descending order of the mean from left to right, and the mean is indicated by an “x”.

These distinct similarities and dissimilarities might also be a reflection of the organs’ primary biological systems and functions. The spleen, thymus, and lymph node are primarily part of the immune system, which could explain their high intra-tissue similarity. Conversely, the large and small intestines mainly serve the digestive system, but they also have immune functions. This dual role might contribute to the distinctive spatial relationships we observed between these two and other three tissues.

For our second approach, we employed the models from leave-one-out cross-validation to predict cell types (see **Methods**) in the held-out images of the same tissue that the model was trained on, as well as images from other tissues. We hypothesized that high predictive accuracy would indicate similar spatial relationships among cell types in the training and prediction images. The prediction accuracy of cell types was quantified using the weighted macro Area-Under-the-Curve (wmAUC, see **Methods**).

The results (Figure 4B) showed high (<0.7) values for all similarities between predicted and original cell types of the same tissue, especially considering the difficulty of predicting a single cell type only from the types of its neighbors. The highest value for comparisons among spleen, thymus, and lymph node were not always those for a tissue with itself; this does not indicate poor performance of the model but rather reflects the similarity between those tissues as already observed above. Those tissues also had a more consistent range of wmAUC values among images from the same tissue compared to those from the small and large intestines. This suggests that regions within the spleen, thymus, and lymph nodes share greater intra-tissue similarity than the intestines.

### Analyzing heterogeneity within tissue images

One assumption of point process models is that point patterns are homogeneous; in our case this means that spatial relationships among cell types remain consistent at different locations within the tissue. However, most tissues have distinct structural and functional units within them (such as stem cell niches). To evaluate whether such organization may be reflected in heterogeneity in cell spatial interaction models, we randomly segmented subregions (tiles) from the original images at two different sizes (5000×5000 and 2500×2500 pixels). Tiles were required to contain at least 100 cells of all cell types and have at least one-fifth of the average number of cells per tile for that image. We ensured that the edges of each tile were at least 500 pixels away from the original image edges, since cells too close to the edge cannot have their interactions accurately counted.

For the same reason, we counted interactions for each cell within a tile with nearby cells outside the tiles. We trained and tested our model on each original image and tile, and for each tile size, we formed a matrix where each row represents a model for a given tile and each column corresponds to a interaction coefficient. Using principal component analysis, we extracted the two major modes of variation, enabling visualization of heterogeneity between individual tile models (Figure 5A-E). We also transformed the interaction coefficients of the model trained on all original images of each tissue using the fitted PCA.

**Figure 5.**
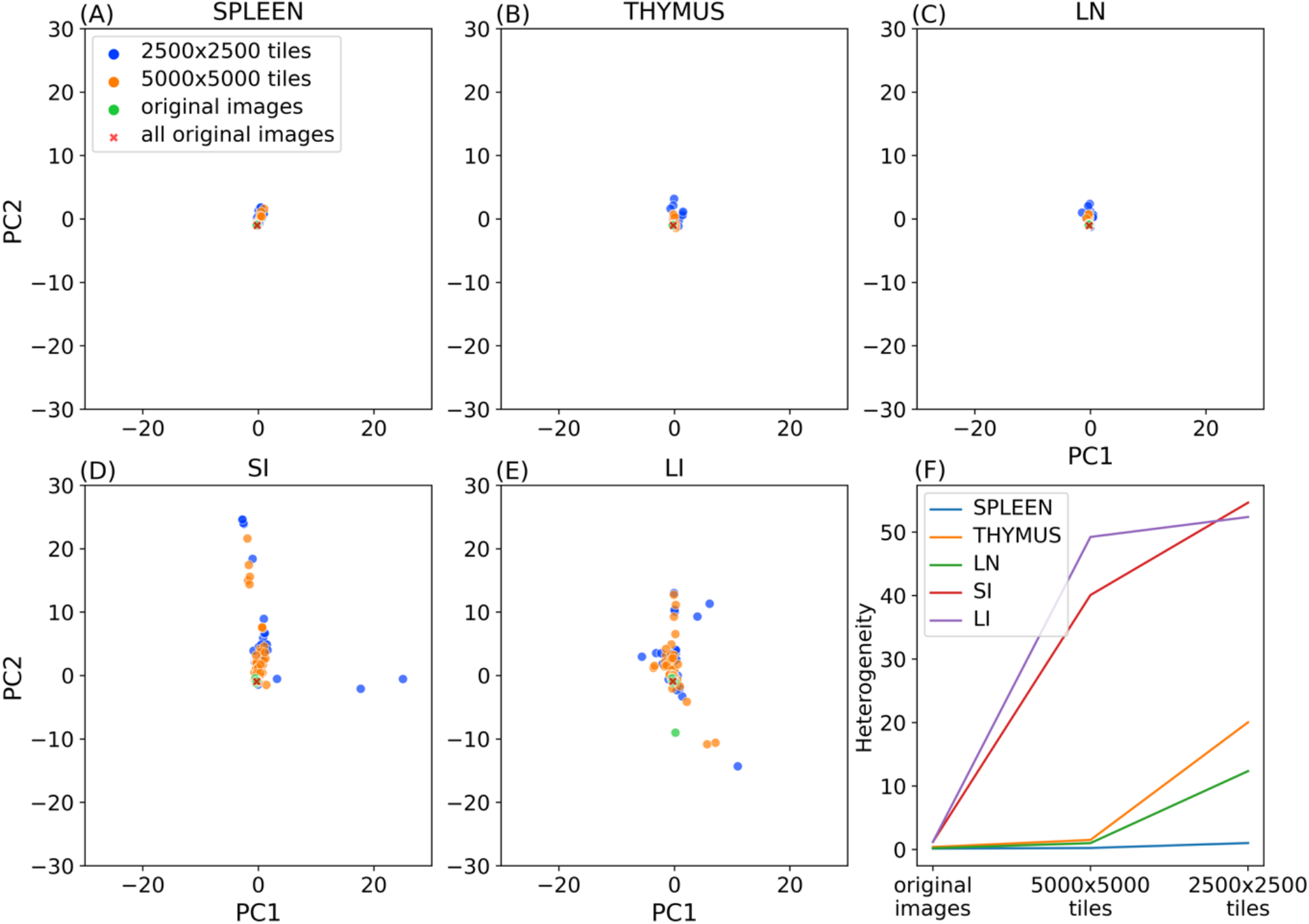
Evaluating tissue heterogeneity of cell type relationships. Panels A to E show the top 2 principal components of the interaction coefficients of various trained models. Panel F illustrates the change of heterogeneity with the tile size for the five tissues.

We also calculated the median of the Euclidean distances between the coefficients of models trained on tiles and coefficients of the model trained on all original images of that tissue. We used this value as a heterogeneity metric (Figure 5F).

As discussed above, spleen, thymus, and lymph nodes displayed lower heterogeneity across their original images compared to those of the large and small intestines. This homogeneity also persists for smaller subregions of those tissues (Figure 5A,B,C) compared to intestine (Figure 5D,E). Figure 5F further quantifies this difference. It is of interest to note that within the three similar tissues, spleen exhibited a much smaller increase in heterogeneity for smaller subregions, suggesting largely homogeneous spatial relationships among cell types across various region sizes in this tissue.

### Visualizing cell type interaction networks

The primary goal of this study was to analyze the spatial relationships among cell types. To summarize our findings, we constructed interaction networks to visualize the interaction coefficients at various ranges in the multirange multitype Strauss Hardcore model (Figure 6).

**Figure 6.**
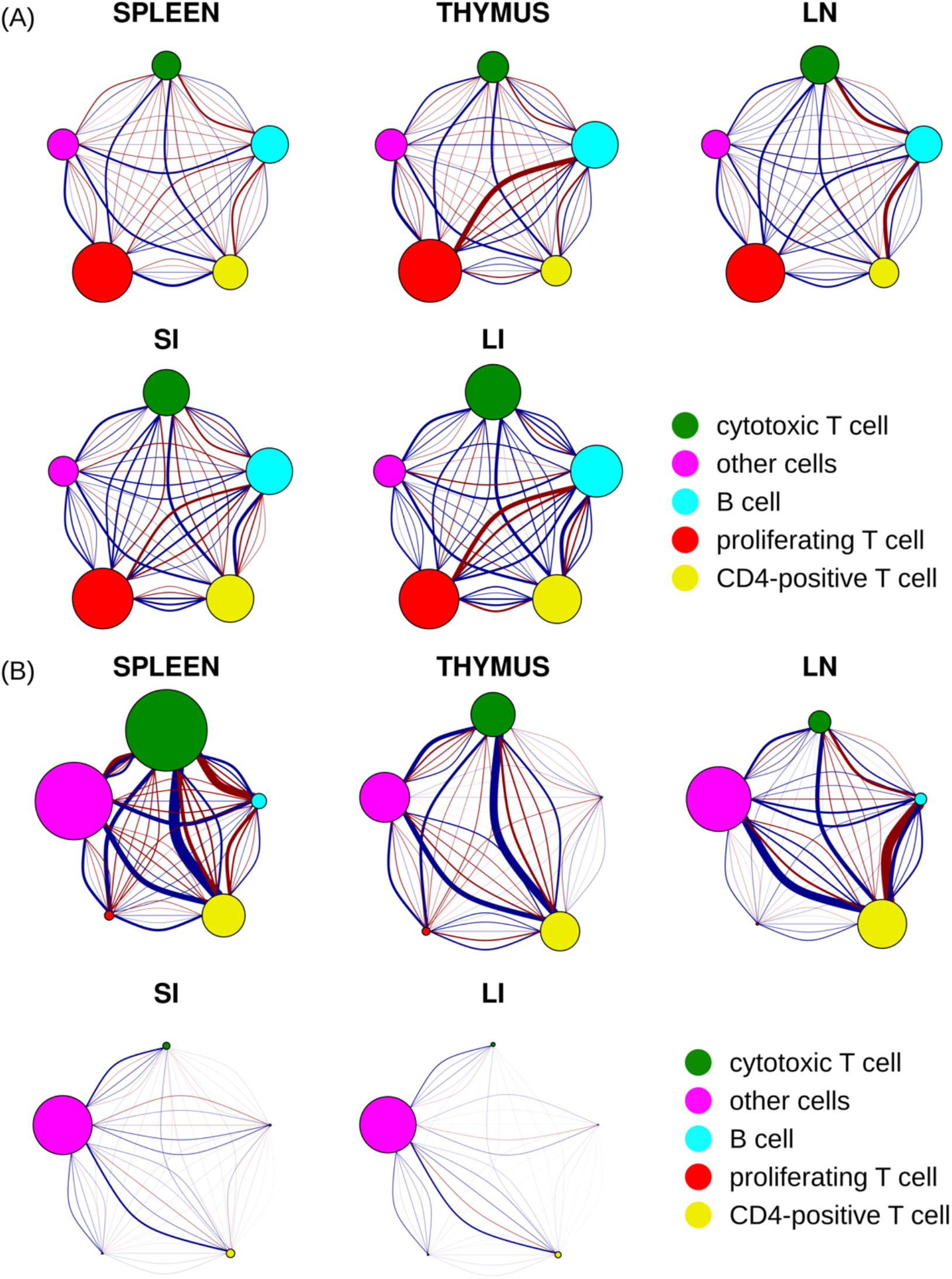
Spatial relationships between five cell types across five different tissues. The size of each node corresponds to the total strength of self-interaction across five distance ranges for that cell type (see Figure S4 for strength of self-interaction at each range). Each pair of nodes is interconnected by five arcs, each representing a different distance range. The range increases from left to right or from bottom to top, with the smallest and farthest ranges corresponding to the most curved arcs. The strength of the relationship between two cell types is depicted by the thickness of the arc, while the nature of their interaction is indicated by the color of the arc (blue as attraction and red as repulsion). (A) A direct, unfiltered illustration by raw interaction coefficients (B) Interaction coefficients adjusted by base intensities of corresponding cell types.

We began by visualizing the interaction coefficients (*δ*) derived from models trained on all images for each tissue type (Figure 6A). These coefficients directly reflect the inherent probability that cell types are near each other, which for simplicity we can interpret as reflecting either “attraction” or “repulsion” between pairs of cell types. However, it’s crucial to emphasize that these inferred interactions aren’t based on isolated pairwise analyses for each pair of cell types. Instead, by integrating the interactions among all cell types in a single point process model, they represent interconnected behaviors between a pair of cell types factoring in influences from all other cell types concurrently.

Our analysis unveiled a variety of noteworthy interaction patterns among different cell types across several tissues. We detected a strong self-attraction among proliferating T cells throughout all the tissues studied (indicated by their larger node diameter). Conversely, cytotoxic T cells and CD4-positive cells demonstrated strong self-attraction in the small and large intestine tissues, but not in the other three tissues. B cells showed moderate self-attraction across all five tissues. As expected, the “other cell” type (cells that could not be annotated given the five markers common to all tissues), exhibited the weakest self-attraction. This is presumably due to the diversity of cell types within this category, with their respective influences offsetting each other.

As also expected, we found that the most intense interactions between two cell types generally occurred within the shortest distance ranges. However, there were a few notable exceptions. The interactions between cytotoxic T cells and B cells in small and large intestine, as well as between proliferating T cells and CD4-positive T cells in the large intestine, were moderate across a range of distances.

Our findings show high consistency between these interaction networks and the analysis presented in Figure 4B. When comparing the interaction networks for the small and large intestines, we discovered high similarity in both the direction of influence (attraction or repulsion) and the intensity of these interactions between cell types, with exception that B cells and proliferating T cells exhibited a notably stronger repulsion against each other within large intestine compared to their counterparts in small intestine. The spleen, thymus, and lymph node also demonstrated a high degree of similarity in terms of the direction of influence (attraction or repulsion) between cell types, but with variance in strength. For instance, thymus displayed stronger repulsion between proliferating T cells and B cells than the other two tissues. Lymph node had a stronger repulsion between B cells and both cytotoxic T and CD4-positive T cells, whereas the spleen demonstrated overall weaker interactions.

Our analysis also highlighted that in spleen, thymus, and lymph node tissues, B cells and CD4-positive T cells displayed a strong repulsive tendency at short distances (less than 40 microns), while they have a moderate attraction at larger distances. Interestingly, the interaction pattern between these two cell types reverses in large and small intestine tissues.

These conclusions are all made by examining the interaction coefficients (*δ*) directly, and thus assumes that the frequencies of the two types are approximately the same. However, it is worth noting that the extent to which a particular interaction is observed in tissue also depends on the base frequencies (β) and the counts (which are also affected by the base frequencies). Therefore, in contrast to “inherent” interaction coefficients presented in Figure 6A, we also calculated “apparent” interaction coefficients by multiplying them with the appropriate base intensities. As shown in Figure 6B, all of the interactions of the “other cells” types were increased across all five tissues after adjustment, due to the high frequency of that type. We found that the self-interaction of cytotoxic T cells in spleen also increased after adjustment. These cells exhibited the strongest repulsion with “other cells” at distances less than 100 pixels (<38 microns) and the strongest attraction at ranges between 100 to 200 pixels (38 to 76 microns). A universal attraction was observed across five tissues between cytotoxic T cells, CD4-positive T cells, and “other cells” with the attraction strength varied. We noted that the repulsion between B cells and both cytotoxic T cells and CD4-positive T cells in the lymph node (LN) persisted after adjustment. Furthermore, all cell types in small and large intestine, excluding “other cells,” displayed minimal self-interaction and minimal interactions among each other after adjustment. This is consistent with the relatively low frequencies of these immune cell types in the small and large intestine tissues.

### Simulating artificial tissue images from generative models

Perhaps the most valuable property of a generative model lies in its ability to create new, realistic data samples based on its learned probability density functions. We therefore asked whether our models could generate artificial tissue images that maintain spatial relationships among cell types.

To begin the simulation process, we generated cell locations using a Poisson distribution that maintained the same total cell density as the original image. We next randomly assigned cell types for all cells based on the density of each cell type in the original image. Following this, we randomly and iteratively selected a cell and reassigned its type according to the cell type counts for that location and the likelihoods derived from the model. This process was continued until the number of sampled cells reached a specified percentage of the total cell count in the image.

We conducted separate trials with different random seeds, and for each trial sampled cells from 0 to 400 percent of total cell counts in intervals of 50 percent. We measured the wmAUC of the original model with respect to the synthetic images, which reflected how well the arrangement of the assigned cell types agreed with the model. We expected that the reassignment process would result in increased wmAUC as it converged as cell type assignments in agreement with the model.

As shown in Figure S1, the wmAUC nearly monotonically increased with the resampling percentage. This observation suggests that our model is capable of generating synthetic images with cell type spatial relationships similar to those in the original images, although the wmAUC values are a bit lower than those obtained for the predicting individual cell types in original images. Even higher accuracy synthetic images could presumably be generated by using even more resampling for different random seeds and choosing the one whose coefficients are most similar to those of the model.

Figure 7 shows how our models can be used to illustrate the differences in cell type arrangement that would result for different tissues if cell locations and sizes were kept constant. To do this, an arrangement of synthetic cells was generated with randomly-chosen cell positions and with cell shapes created from them using a Voronoi diagram truncated at 20 pixels (approximately 7.5 micron radius). Synthetic images were then created from this arrangement using the models trained on all images from each tissue (with 300 percent resampling). The results reflect the trends captured by the adjusted interaction coefficients in Figure 6B for all spatial relationships between cell types, including self-interactions. In particular, the tendency of cytotoxic T cells to be near each other is preserved in all tissues even as the frequency of those cells changes. Cytotoxic T cells and CD4-positive T cells are consistently found near each other across three immune tissues spleen, thymus, and lymph node. This proximity is consistent with their high attraction as represented in Figure 6B. In lymph node synthetic tissue, B cells and CD4-positive T cells exhibit repulsion at short distances whereas attractive to each other at longer distance, aligning with the observations in Figure 6B. While B cells generally appear to be repulsive to both CD4-positive T cells and cytotoxic T cells at short distances in spleen tissue, exceptions can be found Figure 7. This may be attributed to the high intensity of both cytotoxic T cells and CD4-positive T cells in spleen. In both small and large intestine tissues, fewer B cells and T cell types are observed, which is consistent with the low “apparent” interaction strength between these cell types depicted in Figure 6B after adjustment for cell intensity. Nevertheless, we were able to discern the inherent interactions between these cell types in these two tissues, as illustrated in Figure 6A.

**Figure 7.**
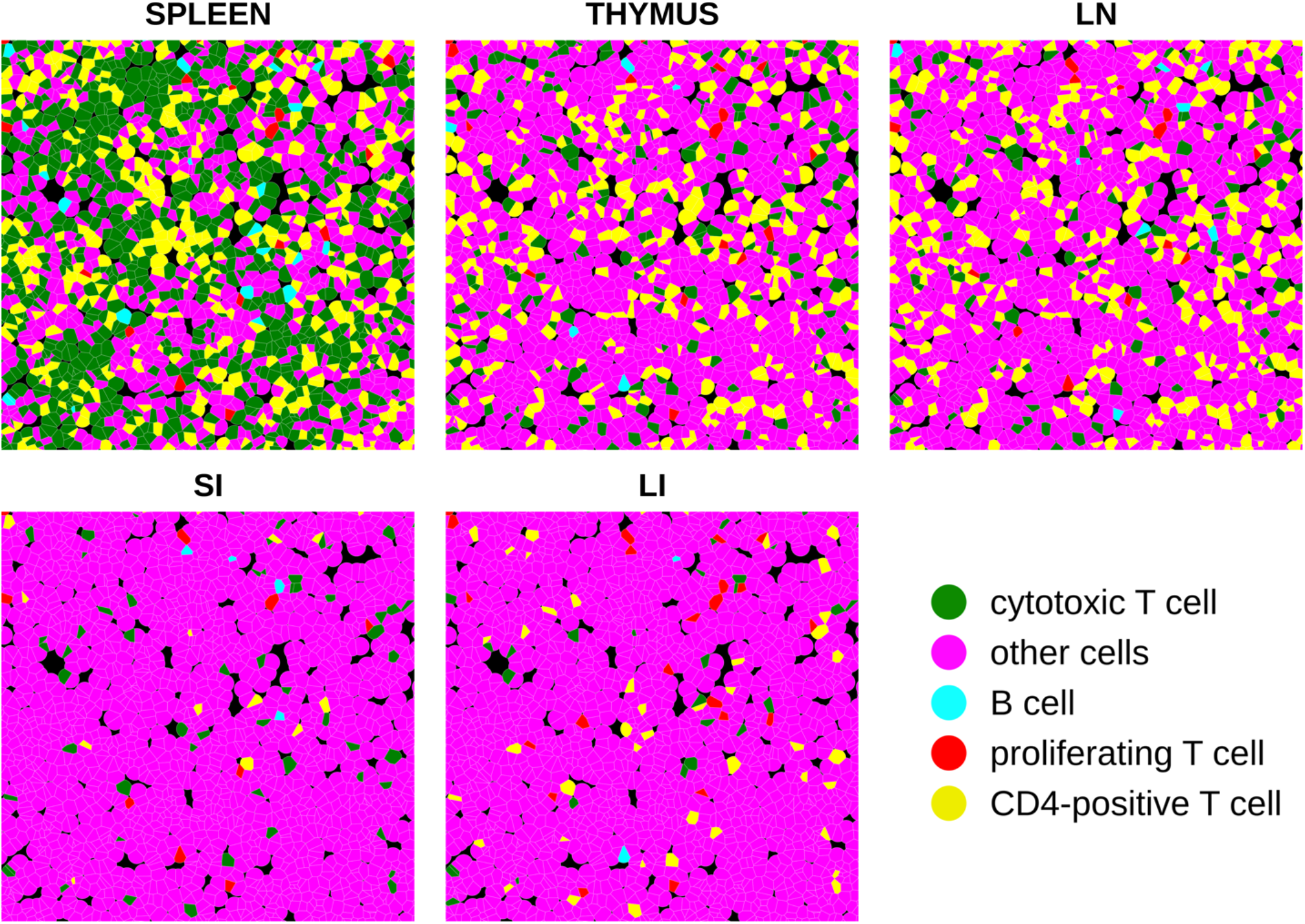
Synthetic tissue images across five tissue types. Each color represents a unique cell type, consistent with representations in other figures.

## Discussion

Spatial relationships among cell types are critical determinants of tissue functions. In this study, we present CytoSpatio – open-source software that constructs innovative generative multitype, multirange point process models to comprehensively learn spatial relationships between 5 cell types in 5 tissues. Our model is built upon a baseline multitype Strauss Hardcore model, incorporating multiple ranges of Strauss radii in a piece-wise manner that captures diverse properties of both signs and strengths of interactions among cell types at varying distances. We demonstrated that our model successfully captures a higher similarity of cell type spatial relationships between images from the same tissue compared to images across different tissues (Figure 4A). Additionally, we provided a quantitative measurement of the spatial heterogeneity within a tissue, revealing the approximate size of heterogeneous structures in five tissues (Figure 5). To visualize the spatial relationships of cell type, we constructed interaction networks and discussed the similarities and differences across 5 tissues (Figure 6). Furthermore, we showcased the capability of our model to generate synthetic tissue images that maintain similar spatial relationships among cell types as those in the original tissue images (Figure 7).

We demonstrated that our multirange, multitype model provides a reasonable approximation for capturing complex spatial relationships among cell types, achieving a balanced trade-off between computational complexity and the ability to learn spatial relationships. In our model, we assumed a maximum range of 500 pixels, or approximately 188 microns, as the distance within which two cells could affect each other. While this is a sound estimation, extending the range further could potentially provide better insight. Furthermore, there is room for refining our model’s interaction function, which currently exhibits a sudden shift of influence every 100 pixels, or approximately 71 microns, due to the piece-wise step function (see Methods). The intervals of our current interaction function could benefit from optimization, and interaction functions with smooth transitions such as Softcore, Fiksel [38], Diggle-Gratton [39], Diggle-Gates-Stibbard [40] might also be worthwhile to explore. In addition, models capturing higher order interactions such as area-interaction [41] and Geyer saturated model [42] where the interaction functions are determined by the relationships of three or more points may be valuable. Currently, the lack of availability of software supporting the multitype versions of the interaction functions limits their use, but future implementations could enhance the representation of interactions among cell types in different scenarios.

Recently, multiplexed tissue imaging technologies have been extended to high-resolution, three-dimensional images [43]. The addition of a third dimension significantly increases the complexity of spatial relationships among cell types and the challenges associated with modeling these relationships. Consequently, there is an urgent need for 3D multitype point process models, since building models on 2D slices or 2D-projections may not capture relationships accurately. We are currently extending our pipeline to model 3D cell type spatial relationships, aiming to deepen our understanding of their impact on tissue function in a 3D context.

Our study successfully depicted the spatial relationships among five cell types in five distinct tissues, with a majority being immune cell types. Rather than making the traditional assumption that these cell types (e.g. B cell, T cell and their subtypes) are generally located near one another for close collaboration [44, 45], we have quantitatively examined their attraction and repulsion tendencies across varying distances. For example, we found a strong preference against B cells and proliferating T cells being closer to each other than ∼38 microns in spleen, thymus, small and large intestine tissues but the opposite tendency at larger distances. Our approach can not only challenge existing qualitative perspectives on spatial relationships among immune cell types but can also potentially provide valuable quantitative insights into how cell types assemble to form tissues.

CytoSpatio effectively simulated cell type locations, accurately reflecting their spatial relationships with one another. We are in the process of upgrading our simulation to include cell shape. To achieve this, we require a generative model capable of learning and simulating diverse cell shapes. In this regard, a robust version of spherical harmonic transform parameterization has been demonstrated as the most effective and accurate method for generating cell shapes [46]. This enhancement will enable us to construct a more comprehensive and detailed representation of tissue images.

## Methods

### Tissue images and cellular data

We used 110 images from the Human BioMolecular Atlas Program (HuBMAP) consortium [37] that had been acquired using the CO-Detection by indEXing (CODEX) [11] method. A summary of these images is provided in Table S1. They were produced by two Tissue Mapping Centers (TMCs): Stanford TMC produced images of the large and small intestine with 47 fluorescence channels (markers), and the University of Florida TMC produced images of the lymph node, thymus, and spleen with 11 fluorescence channels. Image sizes vary, ranging from approximately 5,000 to 15,000 pixels, with each pixel corresponding to a tissue region of 0.37745 × 0.37745 micrometers. The images share five common channels (CD11c, CD21, CD4, CD8, Ki67) across both TMCs. We downloaded files detailing the total intensities of the cell boundary, cytoplasm, nuclear boundary, and nucleus of each channel and the coordinates of cell centers from the HuBMAP portal (https://portal.hubmapconsortium.org/). These files were generated using SPRM (https://github.com/hubmapconsortium/sprm), based on cell segmentations created by Cytokit [47].

### Assigning cell types

Different cell types typically express varying levels of specific cell marker proteins. For instance, proliferating T cells demonstrate high Ki67 levels and low levels of other markers, whereas cytotoxic T cells exhibit high CD8 levels. We defined cell types based only on the five common channels to ensure comparability across tissue types. This decision allows direct comparison of spatial relationships among cell types across various tissues in subsequent analyses.

To compensate for potential differences in channel intensities across tissues, such as those that might arise during image acquisition due to experimental variables like inconsistencies in staining procedures or tissue preparation, we initially z-scored total pixel intensities per cell for each channel within each tissue.

For cell type assignment, we first performed KMeans clustering on the total pixel intensities per cell over the z-scored five common channels across all cells and images from the five tissues. Next, we calculated an overall similarity statistic T based on Gaussian Kernel similarity for intensity compositions of cells between 1) each pair of clusters from KMeans and 2) each cluster from KMeans and each annotated cell type from a lymph node image annotated by Cellar [16] (Figure S2). Using these results as features, we conducted another round of KMeans as meta-clustering to assign the clusters to the five cell types annotated by Cellar.

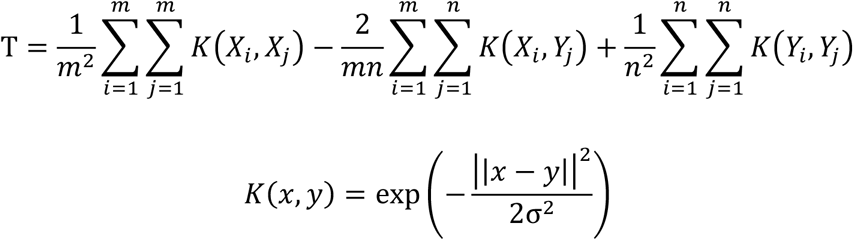

where T is the statistic measuring overall similarity between two cell types, lower T indicates higher similarity. *m* and *n* are the number of cells in two cell types, respectively. *X*_*i*_ and *Y*_*j*_ indicate the cell intensity composition of *i*^th^ cell in cell type *X* and *j*^th^ cell in cell type *Y. K* is the Gaussian kernel similarity and σ is the bandwidth of the kernel (we used 2σ^2^ = 0.08; this value was also used for other Gaussian kernel similarity measurements).

To determine the optimal number of clusters in the initial KMeans, we incrementally increased the number of clusters while monitoring the number of cells in each assigned cell type. We then selected the number of clusters that yielded the highest match between assigned cell types and their corresponding cell types from Cellar (Figure S3). We note that this approach enables the extrapolation of cell type determination from lymph nodes to other tissues, and it allows for finer distinctions within each cell type (i.e., the identification of potential cell subtypes).

For simplicity, all cells assigned to the type “lymphocytes of B lineage” are referred to throughout as simply “B cells.”

### Point pattern and point process model

For each image across 5 tissues, we formed a point pattern ***p*** = {(*x*_1_, *c*_1_), …, (*x*_*i*_, *c*_*i*_), …, (*x*_n_, *c*_n_)}, where *x*_*i*_ is a vector of 2-dimensional coordinates (i.e., cell center) for cell *i, c*_*i*_ is the cell type of cell *i* and *n* is the total number of cells in the image. The coordinates were defined separately in each image. The point patterns belonging to each tissue were considered as random realizations (instances) from a point process model. Our task was to define this point process model.

We assumed cells influence each other by both attraction and repulsion. Therefore, we chose to use the multitype Strauss Hardcore model [21], a kind of multitype Gibbs model, as our baseline model since it satisfies this assumption and can model all cell types at once. The model consists of an expression that allows estimation of the probability density *f*(***p***) of a given point pattern given a set of model parameters (that is, the probability that a particular point pattern would have been observed given those parameters)

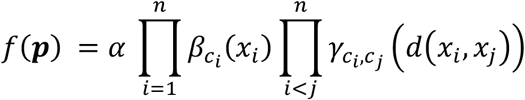

where *f* is the probability density of point pattern ***p***, *α* is a normalizing constant, 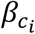 is the intensity of cell type *c*_*i*_ of point *x*_*i*_, *n* is the total number of cells in the pattern, 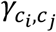 is the interaction function between cell type *c*_*i*_ and *c*_*j*_, *d*(*x*_*i*_, *x*_*j*_), is the Euclidean distance between cell *x*_*i*_ and *x*_*j*_. From this we can also write an expression for the conditional intensity (probability) of finding a cell of cell type *c*_*i*_ at location *x*_*i*_ given the point pattern ***p***

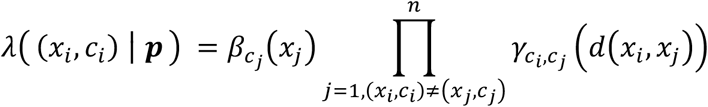

which ignores any contribution from the actual type of that cell.

The interaction function encodes the spatial relationships between two cell types. In multitype Strauss Hardcore model, the interaction function is

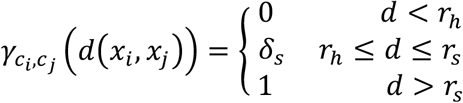

where *r*_h_ is the hardcore radius that stands for the minimum distance that two cells can be from each other, *r*_s_ is the Strauss radius which represents the maximum distance over which cells can affect each other, and *δ*_s_ is the interaction coefficient that captures whether two cells may have attraction (*δ*_s_ > 1) or repulsion (*δ*_s_ < 1) between each other.

One limitation of the conventional Strauss Hardcore model is that the influence between cells is uniformly across a certain single range (Strauss radius *r*_s_), whereas for given spatial relationships between two cell types it may actually vary with distance. To address that, we proposed a multirange multitype model with an upgraded piece-wise interaction function [48]:

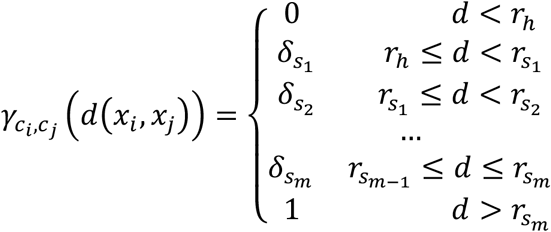

where different interaction coefficients 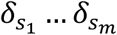 are assigned to each distance interval. For each pair of cell types, we have 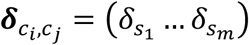, which is the same for all interactions between cell type *c*_*i*_ and *c*_*j*_, where *c*_*i*_, *c*_*j*_ ∈ *C*, and *C* is the set of all cell types.

### Training the point process model

The standard method of fitting point process models to existing data utilizes maximum likelihood estimation (MLE). However, it’s difficult to calculate or approximate the normalizing constant *α* in the probability density function *f* [49]. As an alternative we calculated the log pseudolikelihood:

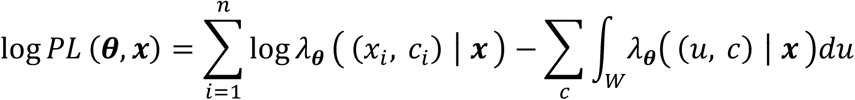

Where ***θ*** = (***β, δ***) is a set of coefficients we need to estimate where 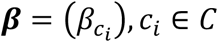 is the first-order term or intensity of each cell type and 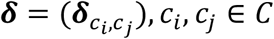 is the set of interaction coefficients between each pair of cell types, *W* is the image window, and the integration is on all possible points *u* over all possible cell types *c* within this window given the point pattern ***x***.

The difficulty of estimating maximum pseudolikelihood is it’s computationally infeasible to integrate over every location within the image window. Therefore, we applied the Berman-Turner quadrature scheme [49, 50] to approximate the background distribution of the conditional intensity function. Each image was evenly split into subregions (tiles) with 20×20 pixels. At the center of each tile and four corners of the image, dummy cells for each of cell types were created. At the location of each real cell, dummy cells for all cell types except the real cell type were also created. This way the integration was converted to a sum weighted by the intensity of cells. The intensity of a cell was calculated by the ratio of the number of cells in its tile to the size of the tile. In other words, cells in the same tile have the same intensity. We therefore had the approximate log pseudolikelihood:

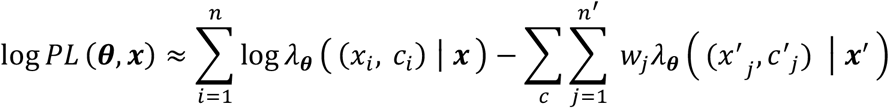

where ***x***^’^ is the new point pattern generated by the quadrature scheme that includes both real and dummy cells, *n*^’^ is the total number of real and dummy cells, and weight *w*_*j*_ is calculated by the area of a quadrature grid (20×20 pixels) over the number of cells in the grid.

We then performed maximum pseudolikelihood estimation by generalized linear model (GLM). The first step was to construct a feature matrix for GLM’s regression (see Table S#). For each point, we converted its cell type to one-hot encoding and counted the number of neighboring cells within a various distance (multirange Strauss radius). The label to predict was the local intensity *y*_i_ = *I*_*i*_/*w*_*i*_, where *I*_*i*_ is an indicator function that equals 1 if current cell is real and 0 if it’s dummy [51, 52].

The whole training process was done by modifying the R package spatstat [53]. We created a new function for our multirange, multitype model.

### Error metric of point process model

Pseudolikelihood can appropriately be used to compare different models trained on the same point pattern. However, pseudolikelihoods for models trained on different patterns are not comparable since those patterns may contain different numbers of cells.

To obtain an error metric that is independent of the training data size, we rewrite the pseudolikelihood as:

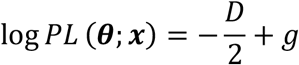

where *g* is a constant and therefore irrelevant in pseudolikelihood comparison. *D* is the deviance that can be written as:

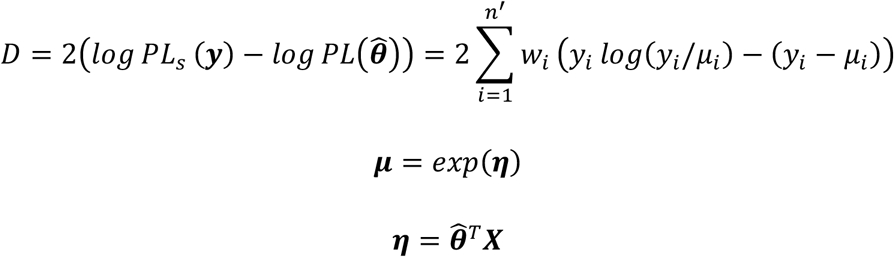

where *log PL*_s_ (***y***) is the log pseudolikelihood of a “saturated” model that has one parameter for each cell to achieve a perfect fit for the data, 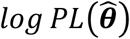, is the log pseudolikelihood of the model under estimation, *w*_*i*_ is weight for cell *i* (definition same as in the equation of log pseudolikelihood), 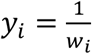 is the true label and ***μ***_*i*_ is the predicted label for cell *i* in GLM. ***X*** is the input feature matrix, 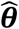 is a vector that contains all base intensity coefficients and interaction coefficients need to be estimated. We assumed the model belongs to exponential family. We therefore applied exponential as the link function of GLM between the linear product ***η*** and predicted label ***μ***.

To account for the influence of data size, we normalized deviance *D* by dividing it by the cell number *n*, yielding the average deviance per cell as our error metric. We interpreted this metric as the average difference between the observed local intensity for each cell and its predicted intensity from a trained model. This metric is particularly sensitive to the value of ***η***. An increase in ***η*** would exponentially elevate ***μ***, leading to a significantly higher average deviance per cell, as exemplified in Figure 2.

### Leave-one-out cross-validation

To prevent overfitting when comparing point process models trained on different tissues, we conducted a leave-one-out cross-validation for each tissue. In this process, we sequentially excluded one image from the current tissue’s training set, fit the model to the remaining images, and predicted the average deviance per cell for the left-out image. As a result, the number of models for each tissue equaled the number of images. In the subsequent analyses, we used them as an ensemble representation of their respective tissues.

### Assessing cell type prediction accuracy

We utilized the Receiver Operating Characteristic (ROC) curve, which is derived from the false positive rate and the true positive rate, to measure the accuracy of cell type prediction. Given that we have five cell types, we need a multi-class ROC; for this, a prediction for one cell type was considered true only if it matched the corresponding cell type and false otherwise.

To calculate overall prediction accuracy, we employed several techniques. First, we calculated the Micro AUC, which considered each cell (independent of its actual type) and counted whether it was correctly predicted. However, a potential issue with Micro AUC arises when class imbalance exists. If a majority of the predictions are biased towards the majority class, Micro AUC could be misleadingly high. This is because the true positive rate and false positive rate in Micro AUC are derived from aggregating predictions across all classes. Consequently, strong performance on the majority class can significantly overshadow any poor performance on the minority classes.

We also computed the Macro AUC to evaluate each cell type independently. This method computes the AUC separately for each class and then averages them, giving equal weight to each class. However, Macro AUC can also be less representative of the model’s overall performance when the class frequencies are different. If a model performs well on a minority class but poorly on a majority class, the Macro AUC might still appear reasonably high despite the model’s overall lower performance on most instances.

We therefore adopted the Weighted Macro AUC (wmAUC) to address this class imbalance issue. Like the Macro AUC, this approach evaluates each cell type independently, but it counters class imbalances by weighting the AUC of each cell type according to its fraction within the total number of cells. Thus, if certain cell types are more common in the dataset, they are assigned more importance in the overall score calculation. Given its effective solution to class imbalance, we chose to use this metric to evaluate the prediction accuracy of cell types.

## Data and code availability

- CytoSpatio software is available at https://github.com/murphygroup/CytoSpatio.
- All data used for this work are available as a reproducible research archive (https://github.com/murphygroup/ChenMurphyCytoSpatioRRA).
- Any additional information required to reanalyze the data reported in this paper is available from the lead contact upon request.

## Acknowledgments

We thank Matthew Ruffalo for helpful discussions. This work was supported in part by grant from the National Institutes of Health Common Fund OT2 OD026682 and OT2 OD033761, and by a traineeship to HC under training grant T32 EB009403.

## Author Contributions

H.C. conceived the study, performed computational analysis, and wrote the manuscript. R.F.M. conceived the study, supervised the study, and wrote the manuscript. All authors approved the content of the manuscript.

