## Supplementary Information for "CytoSpatio: Learning cell type spatial relationships using multirange, multitype point process models"

Computational Biology Department

School of Computer Science

Carnegie Mellon University

Table S1 Summary of tissue images by type and source

| Tissue type | Number of images | Tissue Mapping Center |
|---|---|---|
| Large Intestine (LI) | 26 | Stanford |
| Small Intestine (SI) | 22 | |
| Lymph Node (LN) | 26 | University of Florida |
| Spleen | 24 | |
| Thymus | 12 | |

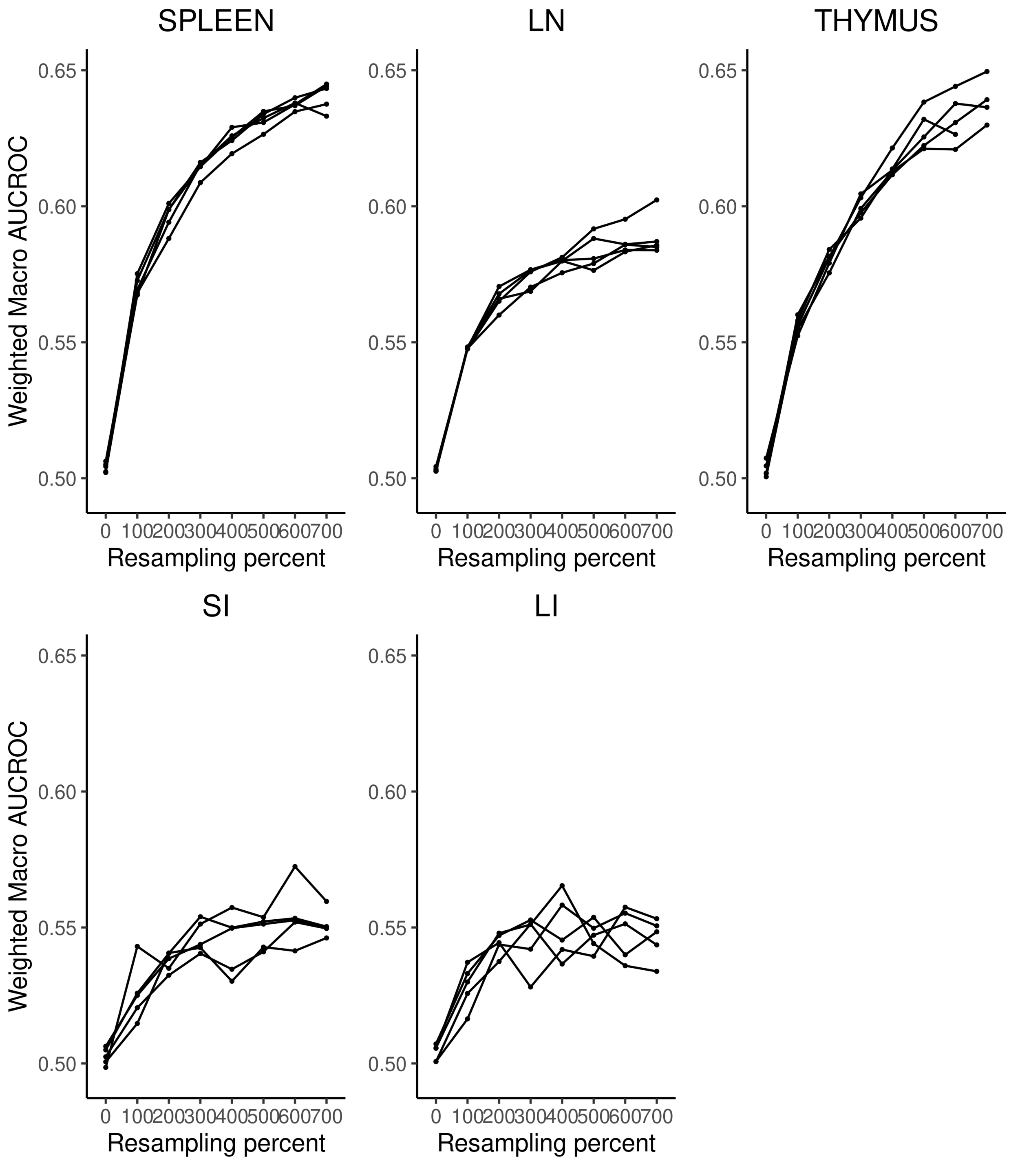


**Figure S1** Evaluation of synthetic tissue image simulation. The weighted macro AUCROC of synthetic images generated using random Poisson cell locations are shown after various amounts of resampling for five tissue types. Each curve plotted corresponds to a synthetic image generated by a model that was trained on an original tissue image. The 'resampling percent' refers to the percentage of the total cell count that were randomly sampled and reassigned according to the model.

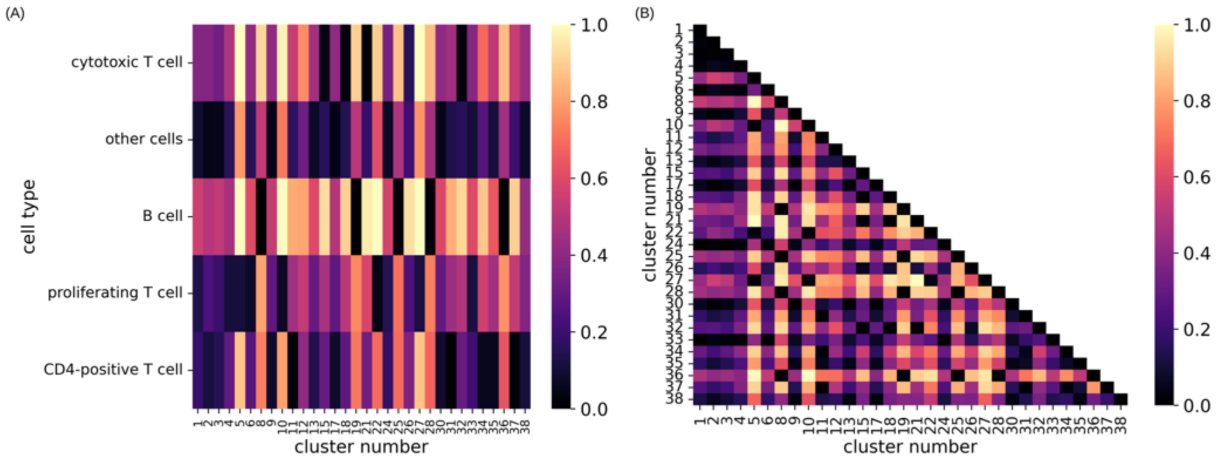


**Figure S2** Gaussian Kernel similarity between cluster centroids visualized for (A) each pair of clusters resulting from KMeans and (B) between each KMeans cluster and each cell type from Cellar. A lighter color indicates higher similarity. KMeans clusters with no cells were excluded from the similarity calculation.

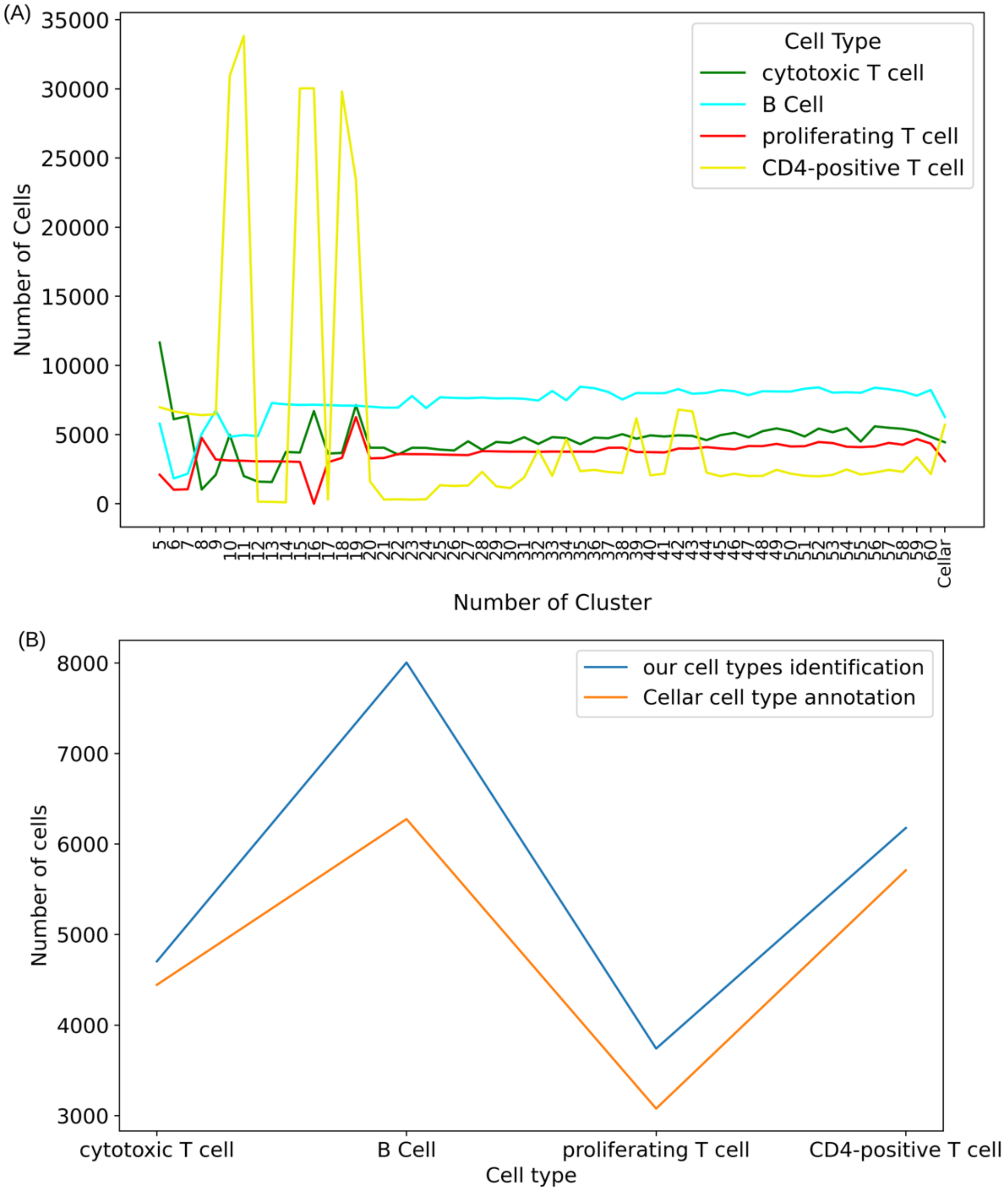


**Figure S3** Defining cell types by comparing cell intensities with Cellar annotations. (A) Determination of the optimal number of clusters in KMeans for cell type definition. The number of clusters was gradually increased until the majority of four Cellar-annotated cell types ("other cells" excluded) showed a consistent cell count. CD4-positive T cells proved the most challenging to identify. We chose 39 as the best cluster number since it presented cell counts most aligned with Cellar annotations, as indicated by the final point on the x-axis. The colors of cell types are consistent with Figure S1. (B) Comparison of our cell type identification and Cellar annotation. Our approach yielded cell counts similar to Cellar annotations with slightly higher numbers for each of cell types. This variation is due to our identification using only 5 shared channels across the five tissue types for cell type classification, in contrast to the 19 channels utilized in Cellar.

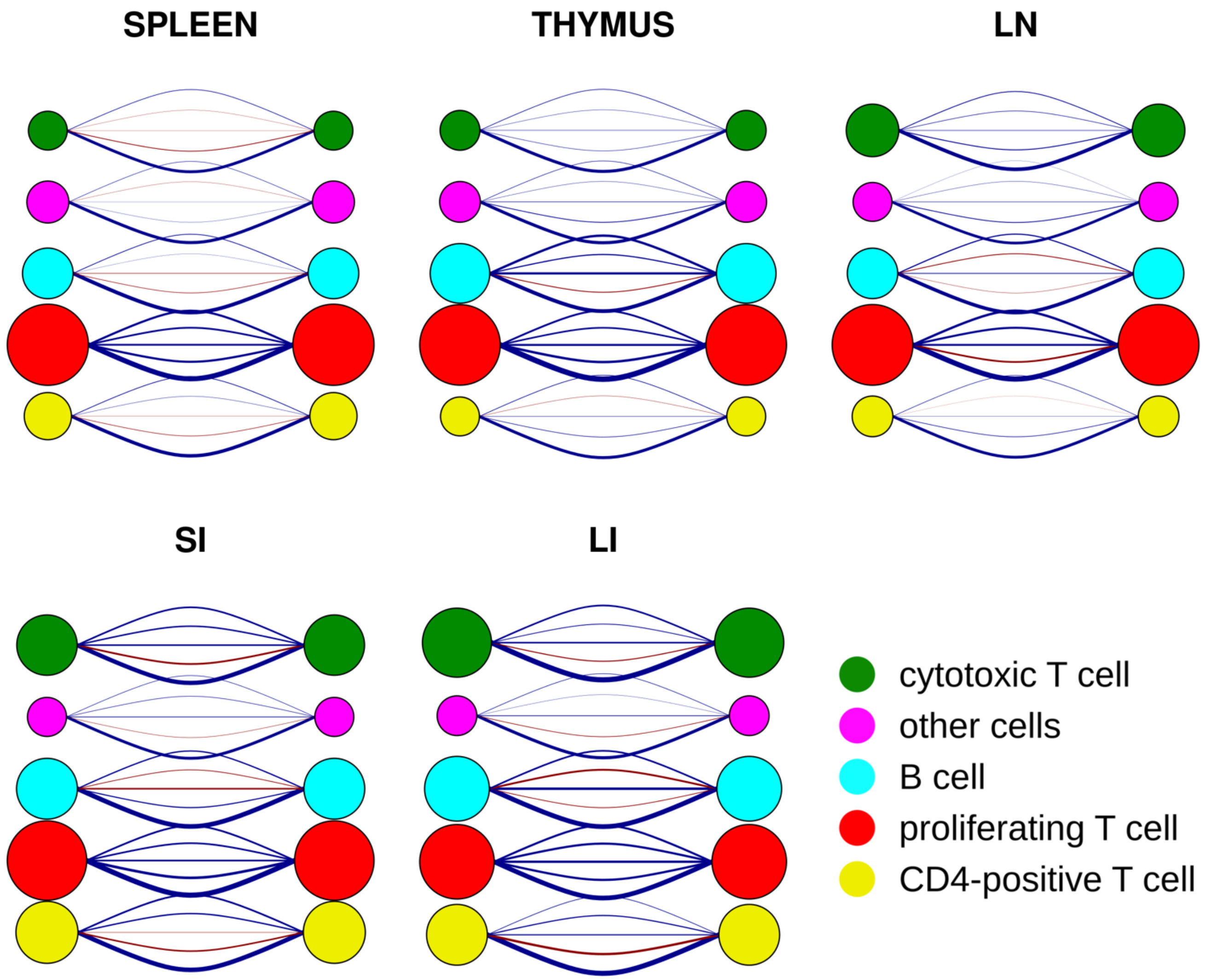


**Figure S4** Self-interactions of five cell types across five different tissues. Each node represents the self-interactions of one cell type. The self-interaction range, which increases from bottom to top, is divided into five arcs. The size of each node corresponds to the total strength of self-interaction for that cell type. The strength of the self-interaction relationship is depicted by the thickness of the arc. The nature of the interaction is indicated by the color of the arc, with blue as attraction and red as repulsion.
